# Overexpression of the *Apoe* gene in the frontal cortex of mice causes sex-dependent changes in learning, attention, and anxiety-like behavior

**DOI:** 10.1101/2024.08.08.607225

**Authors:** Lizbeth Ramos, Abigail E. Harr, Finian L. Zakas, Samuel R. Essig, Griffen J. Kempskie, Nelly A. Fadil, Makayla G. Schmid, Madison D. Pompy, Michael C. Curley, Lisa A. Gabel, Henry L. Hallock

**Affiliations:** Neuroscience Program, Lafayette College, Easton, PA, 18042, USA

## Abstract

Apolipoprotein E (ApoE) is a protein that is important for lipid storage, transport, and metabolism. *APOE* gene variants are associated with Alzheimer’s disease (AD), as well as attentional function in healthy humans. Previous research has shown that *Apoe* transcription is increased following stimulation of the pathway between the locus coeruleus (LC) and frontal cortex (FC) in mice. This result suggests that *Apoe* may affect attentional function by virtue of its expression in circuits that control attention. Does *Apoe* causally regulate attention, or is its expression simply a byproduct of neuronal activity in the LC and FC? To answer this question, we synthetically induced *Apoe* transcription in the FC of male and female mice, and subsequently tested their ability to learn a touchscreen-based rodent version of the continuous performance test of sustained attention (the rCPT). We found that increased *Apoe* transcription impaired performance when attentional demand was increased in male mice, while in female mice, increased *Apoe* transcription significantly accelerated rCPT learning. We further found that this increase in *Apoe* transcription affected subsequent anxiety-like behavior and cellular activity in the FC in a sex-dependent manner. The results of this study provide insight into how *Apoe* causally regulates translationally relevant behaviors in rodent models.

## Introduction

ApoE is a glycoprotein involved in lipid transport and metabolism (Parasuraman et al., 2002). The ApoE protein has two structural domains in humans: The amino-terminal domain (residues 1-191) with receptor and heparin binding regions, and the carboxyl-terminal domain (residues 225-229), which contains the major lipid binding site (Wetterau et al., 1988). There are three predominant isoforms of ApoE in humans (ApoE-e3, ApoE-e2, and ApoE-e4), and each of these isoforms has slight structural differences that affect its function (Zannis et al., 1982). ApoE-e4 is associated with higher risks of heart disease and diabetes, and is linked to increased low-density lipoprotein (LDL) levels and facilitates the conversion of very low-density lipoprotein (VLDL) to LDL, possibly explaining the higher incidence of coronary artery disease in ApoE-e4 carriers (Gregg et al., 1986). ApoE is also highly expressed in the human nervous system (Elshourbagy et al., 1985), where it plays a crucial role in neuroregeneration and degeneration by transporting lipids to axons that are in need of remyelination after nerve damage (Ignatius et al., 1986). ApoE-e4 is also associated with 40-65% of Alzheimer’s disease (AD) cases, and with an earlier onset of AD symptoms (Mahley & Rall, 2000). AD is marked by amyloid beta-protein accumulation in the brain, forming plaques, which are more prevalent in ApoE-e4 carriers following head injury (Nicoll et al., 1996). Neurofibrillary tangles are another AD hallmark, and are caused by tau protein accumulation; while ApoE-e3 binds to tau and may prevent its hyperphosphorylation, ApoE-e4 does not interact with tau, possibly leading to tangle formation (Strittmatter et al., 1994). These differences highlight ApoE-e4’s detrimental role in AD and neurodegeneration, which contrast with ApoE-e3’s potentially protective effects.

The *APOE* gene also contributes to cognitive impairments in both AD patients and the general population, impacting performance on visuospatial attention tasks, such as the cued letter discrimination task, visual search, and vigilance tasks (Greenwood et al., 2000). *APOE* polymorphisms interact with the gene *CHRNA4* to modulate white matter volume and visuospatial attention, with more significant deficits in *APOE*-e4 carriers (Espeseth et al., 2006). Studies using transgenic mice expressing human *APOE* variants also show links between ApoE isoforms and cognition, with ApoE-e3 mice performing better than ApoE-e4 and knockout mice in spatial memory tasks, such as the Morris water maze (Raber et al., 1998). Another study on the combined effect of *APOE* genotype and the pesticide chlorpyrifos on attention and impulsivity showed that ApoE-e4 mice are more compulsive (make more errors of commission) in an attention task, highlighting how *APOE* genotype may affect the brain’s response to injury or toxins and disrupt attention-guided behavior (Peris-Sampedro et al., 2016). *APOE* therefore plays a role in cognition and attention in both patients with neurodegenerative disorders and healthy humans.

Key brain regions involved in attention in humans include the anterior cingulate cortex (ACC) (Carter et al., 1998), and the locus coeruleus (LC) (Aston-Jones et al., 1999). The prelimbic subregion of the frontal cortex (FC) in rodents is anatomically homologous with the primate cingulate cortex (Laubach et al., 2018; van Heukelum et al., 2020), and this region integrates sensory information, plays a role in sustained attention, and monitors competition between conflicting behaviors in both humans and rodents (Carter et al., 1998; Wu et al., 2017). The LC influences arousal and attention, showing increased activity following target stimuli in visual discrimination tasks (Aston-Jones et al., 1999), and sends axonal projections to the FC, where it modulates FC activity during a rodent version of the continuous performance test (rCPT) of attention (Hallock et al., 2024). This task is dependent on FC function (Hvoslef-Eide et al., 2018; Fisher et al., 2020), and drugs that affect catecholamine transmission also affect rCPT performance (Caballero-Puntiverio et al., 2019; 2020), reinforcing the idea that the LC and FC functionally interact to promote attention-guided behavior. A recent study from our lab showed that activation of LC neurons with axonal projections to the FC increased neuronal transcription of the *Apoe* gene in the mouse FC (Craig et al., 2024), suggesting that *Apoe* may affect attention by virtue of its expression in brain areas that are necessary for attention-guided behavior, such as the LC and FC. Despite the known correlations between *Apoe*, attention, and the brain, a causal role for *Apoe* in attention has not been established. Is *Apoe* transcription an epiphenomenon that is merely correlated with circuit function and attention, or does *Apoe* regulate attention? To answer this question, we used a viral targeting strategy to selectively boost *Apoe* transcription in the FC of male and female mice, and subsequently trained these mice on the rCPT. We additionally ran mice on the open field test to illuminate the role of *Apoe* transcription in anxiety-like behavior, and used single-molecule *in situ* hybridization (RNAscope) to correlate individual differences in *Apoe* transcription with behavior and cellular function in the FC.

## Results

On average, the number of sessions needed to reach performance criteria on stage 1 (Fig. 1d), stage 2 (Fig. 1e), and stage 3 (Fig. 1f) did not significantly differ between either sex or group (*p* > 0.05 for group x sex two-way ANOVAs). Out of all stages, mice reached performance criteria on stage 2 most quickly, and took the longest to reach performance criteria on stage 3. During the first five stage 3 training sessions, hit latencies increased for both groups in female (*F*(1,4) = 2.558, *p* = 0.0477, main effect of session; Fig. 2a), but not male (Fig. 2b), mice, suggesting that hit reaction times during learning in the rCPT are sex-dependent. In contrast, no statistically significant differences in false alarm latencies were observed as a function of session or group in female mice (Fig. 2c), but a significant group x session interaction was observed between the final stage 3 and probe sessions in male mice (*F*(1,1) = 4.278, *p* = 0.0496; Fig. 2d), with false alarm latency being lower in male Apoe mice during the final stage 3 session compared to EGFP controls, but evening out during the probe session. No statistically significant differences in reward latency were observed as a function of session or group for female mice (although an outlier resulted in large error bars during the final stage 3 and probe sessions; Fig. 2e), or male mice (Fig. 2f).

**Fig. 1:**
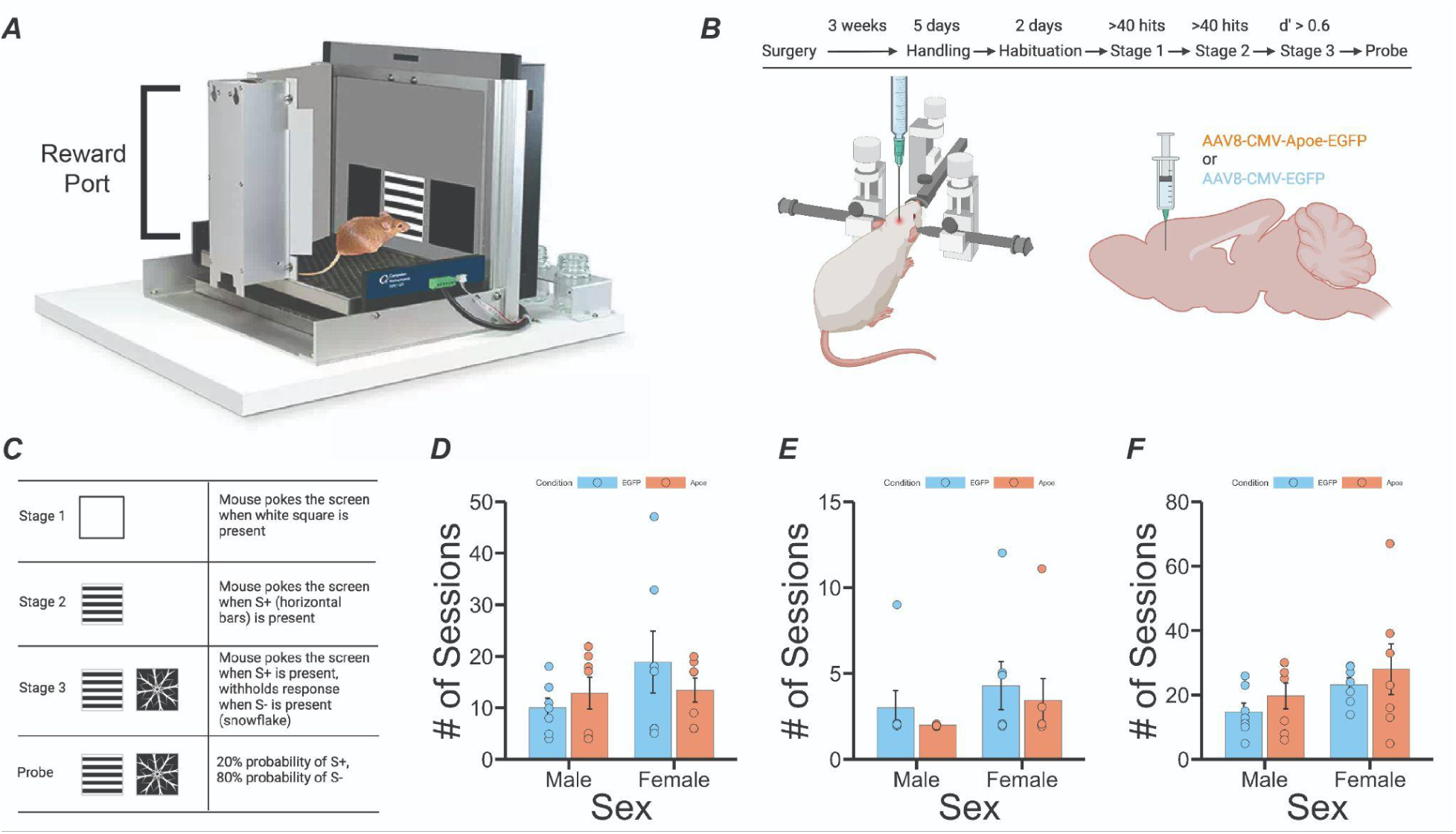
Experimental design and number of sessions needed to reach learning criteria for the rCPT. *(A)* Chamber design. *(B)* Experimental design and surgery schematic. *(C)* Description of rCPT stages. *(D)* There were no significant differences in the number of sessions needed to reach criteria on stage 1 between group (Apoe vs. EGFP controls) or sex. *(E)* There were no significant differences in the number of sessions needed to reach criteria on stage 2 between group or sex. *(F)* There were no significant differences in the number of sessions needed to reach criteria on stage 3 between group or sex.

**Fig. 2:**
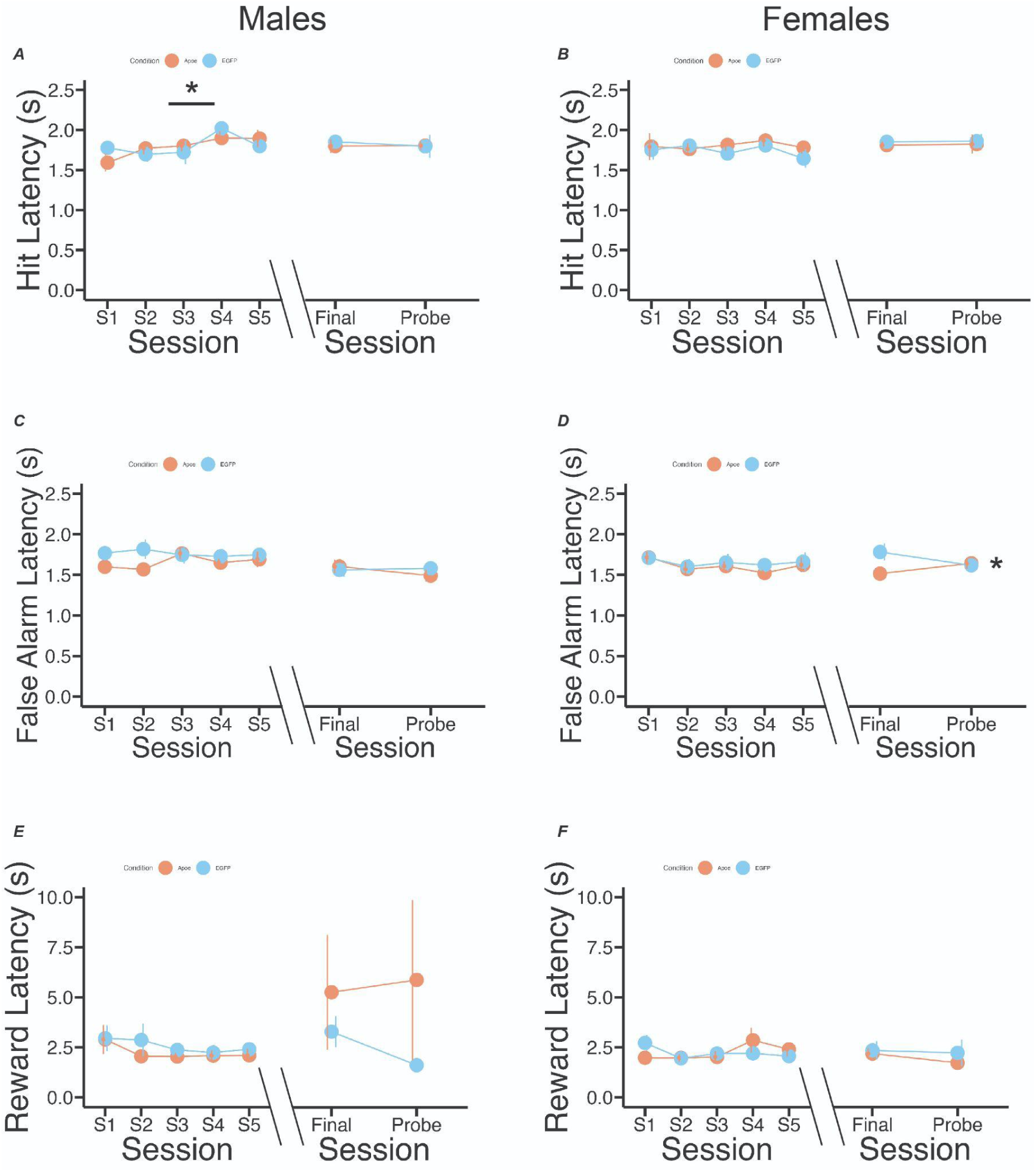
Latencies during rCPT learning and performance. *(A)* Hit latency increases across the first five sessions for both groups in females (*F*(1,4) = 2.558, *p* = 0.0477, main effect of session), but does not significantly differ between groups. No significant differences in hit latency were observed between groups or between the final stage 3 and probe sessions in females. *(B)* No significant differences in hit latency between sessions or groups was observed for males. *(C)* No significant differences in false alarm latency between sessions or groups was observed for females. *(D)* No significant differences in false alarm latency between sessions or groups was observed during rCPT learning for males, but a significant group x session interaction was observed for males between the final stage 3 and probe sessions (*F*(1,1) = 4.278, *p* = 0.0496). *(E)* No significant differences in reward latency between sessions or groups was observed for females, or *(F)* males.

In female mice, hit rate was significantly higher in Apoe mice compared to EGFP mice during the first five stage 3 sessions (*F*(1,1) = 7.689, *p* = 0.00739, main effect of group; Fig. 3a), but did not differ significantly between groups in male mice (Fig. 3b). These results suggest that *Apoe* over-expression in the FC of female mice increases their ability to detect an S+ during learning, without altering their reaction times during hits. False alarm rates (the mouse’s ability to detect an S-) did not significantly differ between groups in either male (Fig. 3c), or female (Fig. 3d), mice. Discrimination index, or d’, scores were significantly higher during the first five stage 3 training sessions (*F*(1,1) = 11.308, *p* = 0.00135, main effect of group), but not during the final stage 3 or probe session, in female Apoe mice (Fig. 3e). In male mice, d’ scores significantly increased across the first five stage 3 sessions in both groups (*F*(1,4) = 3.472, *p* = 0.0129, main effect of session), and d’ scores were significantly lower in the Apoe group compared to the EGFP group in the final stage 3 and probe sessions (*F*(1,1) = 9.881, *p* = 0.0044, main effect of group; Fig. 3f), indicating that *Apoe* over-expression in the FC improves rCPT learning in female mice, but reduces rCPT performance in male mice that have already learned the task. To assess whether differences in d’ score were due to mice responding more conservatively (withholding responses more), or more liberally (responding more often), we calculated a c score, which is defined as the sum of the z-scored hit and false alarm rates divided by two. A higher c score represents a more conservative response bias, while a lower c score represents a more liberal response bias (Stanislov and Todorov, 1999). In female mice, c scores were significantly lower in the Apoe group during the first five stage 3 sessions (*F*(1,1) = 4.902, *p* = 0.0306, main effect of group; Fig. 3g). In contrast, c score did not significantly differ between groups in male mice (Fig. 3h).

**Fig. 3:**
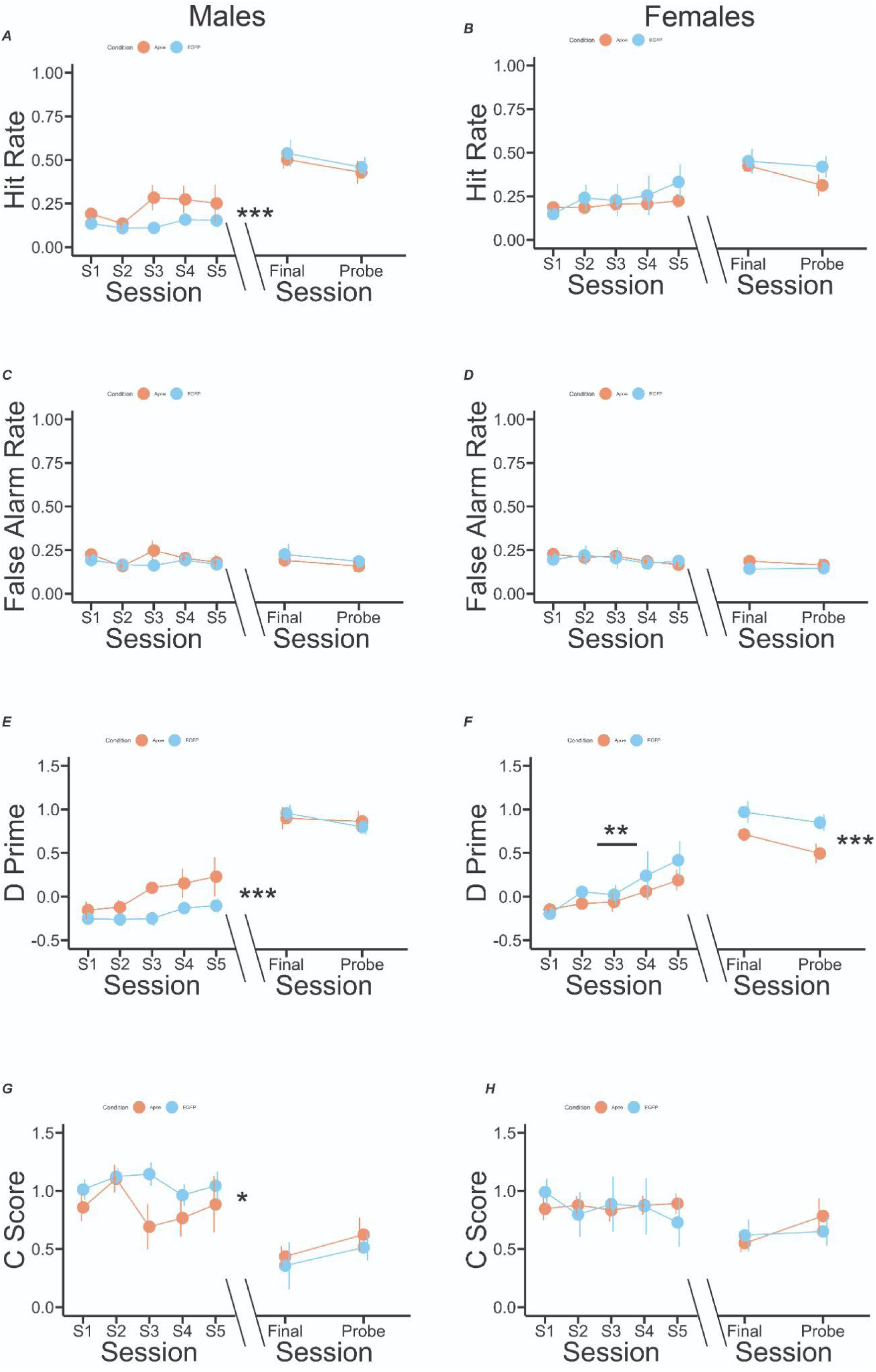
Performance metrics for rCPT learning and performance. *(A)* Hit rate significantly differed between groups during rCPT learning in females (*F*(1,1) = 7.689, *p* = 0.00739, main effect of group); in contrast, hit rate did not significantly differ between session or group during asymptotic stage 3 performance and the probe session in females. *(B)* Hit rate did not differ significantly between sessions or groups during rCPT learning or asymptotic performance in males. *(C)* False alarm rate did not differ significantly between sessions or groups during rCPT learning or asymptotic performance in females, or *(D)* males. *(E)* D’ was significantly higher in Apoe mice compared to controls during rCPT learning in females (*F*(1,1) = 11.308, *p* = 0.00135, main effect of group), but not between the final stage 3 and probe sessions. *(F)* D’ significantly increased across sessions during rCPT learning in males, regardless of group (*F*(1,4) = 3.472, *p* = 0.0129, main effect of session), but was significantly lower in Apoe mice compared to controls during the final stage 3 and probe sessions (*F*(1,1) = 9.881, *p* = 0.0044, main effect of group). *(G)* C score was significantly lower in Apoe mice compared to controls during rCPT learning in females, indicating a more liberal response bias (*F*(1,1) = 4.902, *p* = 0.0306, main effect of group), whereas c score did not significantly differ between groups during the final stage 3 and probe sessions in females. *(H)* C score did not differ significantly between sessions or groups during rCPT learning or asymptotic performance in males.

To assess whether differences in rCPT performance could be due to general differences in anxiety-like behavior or motor control, we ran mice on the open field test following completion of the rCPT (Fig. 4a), and measured the number of entries made into the center of the arena, time spent in the center of the arena, time spent freezing in the arena, and total distance traveled during the testing session. We observed a group x sex interaction for number of center entries, such that female Apoe mice made less center entries than female EGFP mice, but male Apoe mice made more center entries than male EGFP mice (*F*(1,4) = 7.581, *p* = 0.0126; Fig. 4b), suggesting that *Apoe* over-expression in the FC may have sex-dependent effects on anxiety-like behavior. We did not observe any statistically significant differences between sexes or groups for the amount of time spent in the center of the arena (Fig. 4c), time spent freezing (Fig. 4d), or total distance traveled (Fig. 4e).

**Fig. 4:**
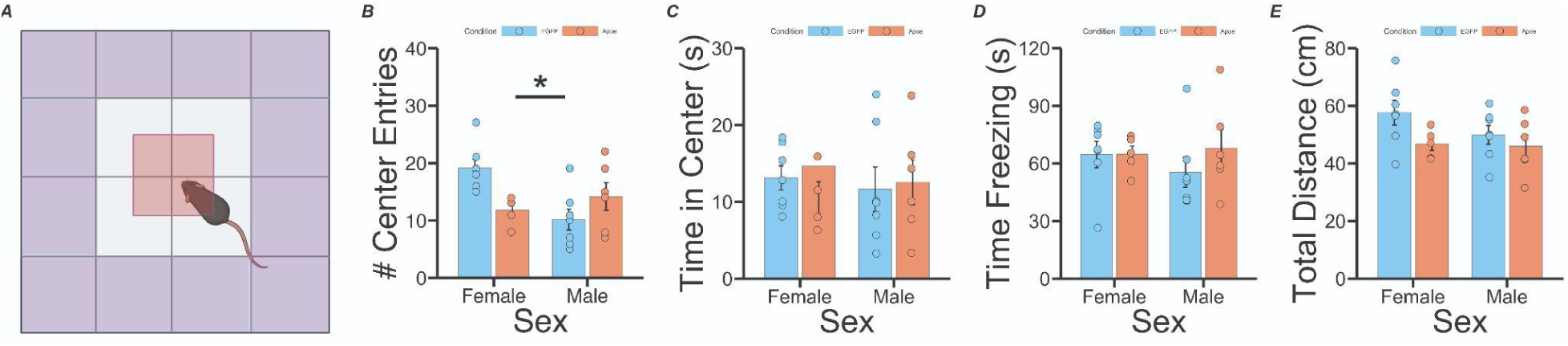
Open field data. *(A)* Schematic of open field test, with outer edge (purple) and center (red) highlighted. *(B)* A significant group x sex interaction was observed for the number of center entries, with female Apoe mice making less center entries than controls, and male Apoe mice making more center entries than controls (*F*(1,4) = 7.581, *p* = 0.0126). *(C)* There were no statistically significant differences in the number of time spent in the center, *(D)* time spent freezing, or *(E)* total distance traveled as a function of group or biological sex.

In order to ensure that our viral strategy boosted levels of *Apoe* transcription in the FC, we killed all mice following open field testing, and used single-molecule *in situ* hybridization to visualize virally-derived *Gfp*, *Apoe*, and *Fos*, which is an immediate early gene (IEG) commonly used as a marker of neuronal activity (Fig. 5a). Transcripts from each gene were readily visible in coronal slices of FC tissue (Fig. 5b). We found that the proportion of DAPI-stained nuclei that co-expressed *Gfp*+ was not significantly different between groups or sexes (Fig. 5c), demonstrating that the viruses infected a similar number of cells. We also observed that the number of *Apoe* transcripts per cell (DAPI-stained nucleus) was significantly higher in Apoe mice (*F*(1,1) = 13.256, *p* = 0.00174, main effect of group; Fig. 5d), as was the number of *Apoe* transcripts per *Gfp*+ cell (*F*(1,1) = 8.178, *p* = 0.01, main effect of group; Fig. 5e), indicating that our viral overexpression strategy did indeed boost levels of *Apoe* transcription in the experimental group. In order to link *Apoe* expression with cellular activity more generally, and probe whether *Apoe* transcription is associated with active neurons, we also measured the number of *Apoe* transcripts per *Fos*+ cell, and found a group x sex interaction, such that number of *Apoe* transcripts per *Fos*+ cell was significantly lower only in male Apoe mice (*F*(1,1) = 4.367, *p* = 0.05; Fig. 5f). To probe whether individual differences in *Apoe* transcription are associated with fluctuations in *Apoe* transcription in *Fos*+ cells specifically, we correlated number of *Apoe* transcripts per cell with number of *Apoe* transcripts per *Fos*+ cell, and found that this relationship differed between male and female mice, with a positive correlation observed in female mice, and a negative correlation observed in male mice (Cohen’s q = 0.792, *p* = 0.047, Fisher’s z-test; Fig. 5g). Taken together, these results show that the relationship between *Apoe* transcription and cellular activity in a local cortical network differ between biological sexes, which may partially explain sex-dependent differences in attention-guided and anxiety-like behavior between male and female mice in the current study. To investigate whether individual differences in *Apoe* expression are linked with anxiety-like behavior specifically, we correlated number of *Apoe* transcripts per cell with number of center entries in the open field test, and found again that this relationship differed between male and female mice, such that these variables were negatively correlated in female mice, and positively correlated in male mice (Cohen’s q = −0.872, *p* = 0.032, Fisher’s z-test; Fig. 5h). Correlations between *Apoe* transcription and rCPT metrics (d’, c score, hit and false alarm rates, and latencies) did not reveal any statistically significant relationships.

**Fig. 5:**
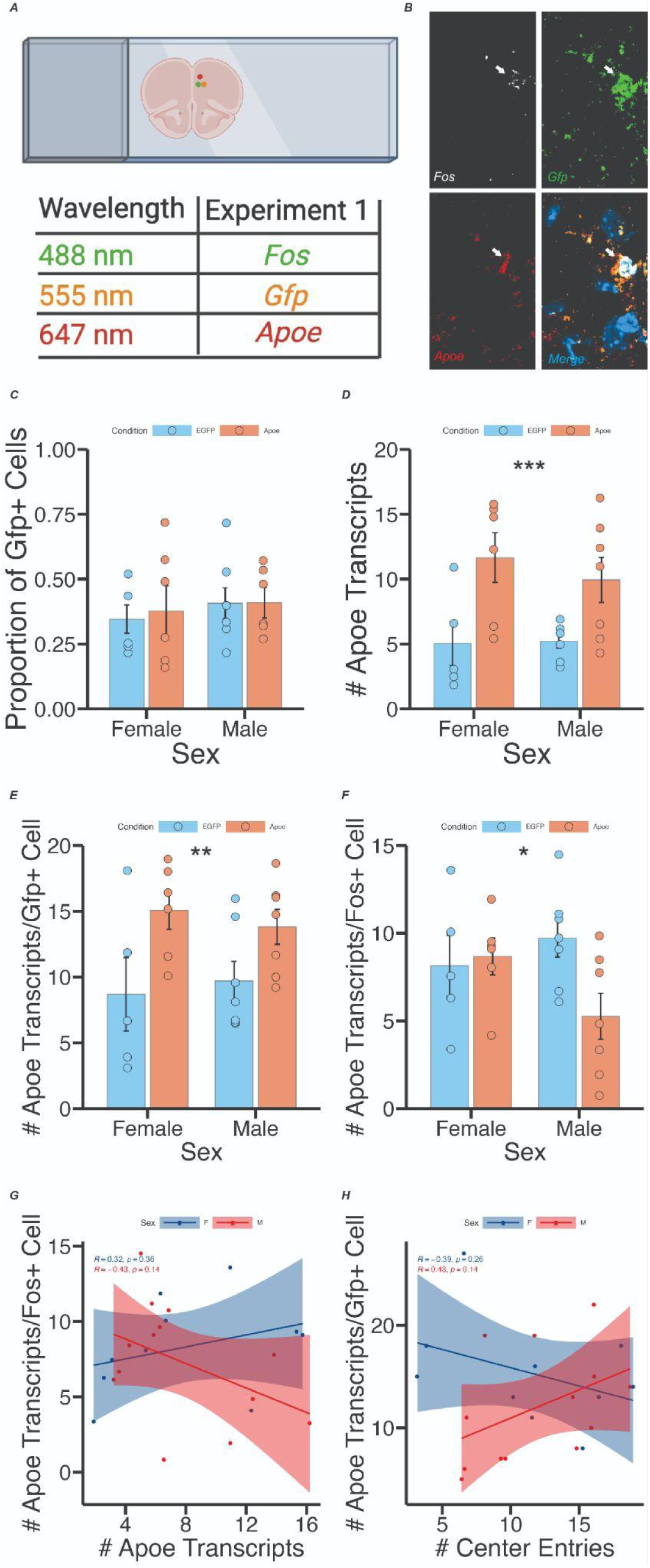
*Apoe* expression in the FC. *(A)* Schematic of experimental design for RNAscope verification of *Apoe* expression in the FC. *(B)* Representative confocal image showing co-expression of *Fos*, virally-derived *Gfp*, and *Apoe* in FC tissue. *(C)* No significant differences in proportion of cells expressing *Gfp* were observed as a function of group or biological sex. *(D)* The average number of *Apoe* transcripts per cell was significantly higher in the Apoe group compared to controls in both sexes (*F*(1,1) = 13.256, *p* = 0.00174, main effect of group). *(E)* The average number of *Apoe* transcripts per *Gfp*+ cell was significantly higher in the Apoe group compared to controls in both sexes (*F*(1,1) = 8.178, *p* = 0.01, main effect of group). *(F)* A group x sex interaction was observed for average number of *Apoe* transcripts per *Fos*+ cell, such that the number of *Apoe* transcripts in *Fos*+ cells was lower in male Apoe mice (*F*(1,1) = 4.367, *p* = 0.05). *(G)* Correlations between the number of *Apoe* transcripts per *Fos*+ cell, and the number of *Apoe* transcripts per cell, were significantly different between male and female mice (Cohen’s q = 0.792, *p* = 0.047, Fisher’s z-test). *(H)* Correlations between the number of *Apoe* transcripts per cell, and the number of center entries in the open field test, were significantly different between male and female mice (Cohen’s q = −0.872, *p* = 0.032, Fisher’s z-test).

## Discussion

We find that boosting levels of *Apoe* transcription in the FC has sex-dependent effects on learning, sustained attention, anxiety-like behavior, and cellular function. Specifically, female Apoe mice had higher levels of performance during initial rCPT training (measured by an increase in d’ score) compared to controls. This performance increase was driven by a higher hit rate and a more liberal response bias. In contrast, *Apoe* overexpression in the FC did not cause a significant alteration in d’, hit rate, or response bias during initial rCPT sessions in male mice, indicating that *Apoe* expression in the FC affects task learning in a sex-dependent manner. Notably, we observed sex differences in rCPT learning at baseline; male control mice had higher overall d’ prime scores during the initial five sessions than female control mice, which was driven by correspondingly lower c scores in males, indicating that male mice may learn the rCPT more quickly than females because they respond less conservatively. Female Apoe mice had higher d’ prime scores that were driven by lower c scores, suggesting that *Apoe* overexpression effectively made female mice adopt a “male-like” strategy during initial rCPT training. To our knowledge, only one other study has done a direct comparison between male and female mice during rCPT learning (DeBrosse et al., 2022), and this study reported that female mice had lower d’ scores than male mice during initial training, but that this difference was driven by lower c scores in females. The reason for the discrepancy between our findings and the findings of this study are unclear, but studies with human CPTs consistently find that females have lower d’ scores and higher c scores (more conservative response biases) than males (Connors et al., 2003; Burton et al., 2010), which are in line with our results. We further found that d’ scores were lower in overtrained male Apoe mice during the final stage 3 and probe sessions, indicating that *Apoe* overexpression did not affect learning rate, but rather limited the ability of male mice to discriminate between the S+ and S-(lowered the “ceiling” for proficiency on the task) - this effect was not observed in female mice. This result indicates that *Apoe* overexpression causes a decrease in sustained attention in males, with male Apoe mice performing well enough to pass performance criteria for stage 3, but never achieving the same level of mastery on the task compared to controls. One possible explanation for the lack of effect in females is that increased performance early in training in female Apoe mice may mask attention deficits once performance criteria are met. Future studies could disentangle effects of *Apoe* expression on learning and performance in the rCPT by injecting the virus at distinct time points during training (in naive vs. overtrained mice, for example).

Lesions of the FC reliably disrupt rCPT performance in male rats (Fisher et al., 2020), and lesions of a smaller sub-region of the FC increase false alarm rates and lower c scores in male mice (Hvoslef-Eide et al., 2018), corroborating the idea that the FC is necessary for the performance of this task. Our findings suggest, therefore, that *Apoe* overexpression affects cellular function in the FC in opposite ways in male and female mice. Human studies also show links between activation in the cingulate cortex and CPT performance (Ogg et al., 2008; Tana et al., 2010), highlighting the translational validity of studying this brain region in the CPT across species. Interestingly, we observed that the number of *Apoe* transcripts per *Fos*+ cell was lower in male Apoe mice, but unaffected in female Apoe mice, indicating that *Apoe* may have different effects on cellular excitability in local networks that are sex-dependent. A recent paper from our lab showed that *Apoe* expression in the FC following stimulation of the locus coeruleus is highest in somatostatin-expressing interneurons in female, but not male, mice (Craig et al., 2024). Differences in brain function and behavior following *Apoe* overexpression in our study could therefore be due to differences in cell type-specific expression of *Apoe* between sexes. We used a cytomegalovirus (CMV) promoter to drive Apoe and EGFP expression in our mice, which has pan-neuronal tropism, as well as weak tropism for glial cells in rodents (Yaguchi et al., 2013). Future studies using this approach should characterize which cell types (glia, excitatory neurons, inhibitory neurons) predominantly express virally-derived *Apoe* in order to dissect the mechanisms by which *Apoe* affects brain function and behavior.

In order to assess whether performance differences in the rCPT could be due to general differences in motor behavior or anxiety, we tested all mice in the open field following successful completion of the rCPT probe session. We found no differences in total distance traveled as a function of group or biological sex, suggesting that differences in attention-guided behavior in Apoe mice were not due to general alterations in locomotion. We also did not find any differences in the amount of time spent freezing, or time spent in the center of the open field arena. We did, however, find an effect of *Apoe* overexpression on number of center entries, and this effect was again sex-dependent, such that female Apoe mice made less center entries (displayed higher anxiety-like behavior), and male Apoe mice made more center entries (displayed lower anxiety-like behavior) compared to within-sex controls. Differences in anxiety-like behavior may therefore partially underlie differences in attention-guided behavior observed between groups in the rCPT. Interestingly, female Apoe mice had higher hit rates, a less conservative response bias, and higher d’ scores during rCPT learning (indicating a higher willingness to respond, and potentially less anxiety in the touchscreen chambers), which stands in contrast to the reduced number of center entries observed in female Apoe mice during the open field test. One possible explanation for these disparate results is that effects of *Apoe* overexpression on behavior are context-dependent, and that any effects on rCPT performance are unrelated to effects on open field behavior. Another possible explanation is that the number of center entries may not, in and of itself, be sufficient to identify an anxiety-like phenotype, as other indicators of anxiety-like behavior (freezing and time spent in center) were not significantly different between group or biological sex. Metrics of rCPT performance (hit rate, false alarm rate, d’, and c score) were not correlated with the number of center entries for either sex, providing evidence that behavior on the open field test and behavior during the rCPT are separate phenomena. These results may differ from those in humans, as at least one study has demonstrated a positive correlation between chronic anxiety and problem solving skills (Wisconsin card sorting task performance) in cognitively normal human *APOE*-e4 carriers (Casselli et al., 2004); however, other studies have shown a negative correlation between anxiety and cognition in *APOE*-e4 carriers (Stonnington et al., 2011), highlighting the complicated relationship between *APOE* and behavior. An additional study reports improvements in spatial memory that are accompanied by increases in anxiety-like behavior in female mice expressing the human ApoE-e4 protein (Siegel et al., 2012), supporting the hypothesis that Apoe expression can, in some instances, increase both cognitive performance and anxiety concurrently. Several studies have reported general elevations in anxiety in male ApoE-e4 carriers (Robertson et al., 2005; Holmes et al., 2016), as well as male mice expressing human ApoE-e4 (Raber, 2007), with mice in the latter study showing alterations in amygdala function. Given that the FC is also highly involved in anxiety-like behavior in rodents (Lacroix et al., 2000; Suzuki et al., 2016; Elliott et al., 2016), it follows that *Apoe* expression in this brain area would also affect these behaviors.

Although the relationships between *APOE* genotype and attention in humans is well-established (Greenwood et al., 2000; Parasuraman et al., 2002), it is unclear how *APOE* affects brain regions that are important for attention. Furthermore, there remains a paucity of studies examining sex-specific effects of *APOE* and attention in both humans and mice; such studies are critically important in light of potential differences in vulnerability for Alzheimer’s disease (AD) between males and females (Mielke et al., 2014; Snyder et al., 2016; Nebel et al., 2018). In the current study, we attempt to provide a causal link between *Apoe* expression, brain function, and attention-guided and anxiety-like behaviors. Notably, the majority of studies in both humans and mice that study ApoE have focused on *APOE* alleles, as these are highly relevant for human health. It is difficult to compare the results of studies that use transgenic mice expressing a human allele with the results of the current study for several reasons. First, the human *APOE* gene is structurally distinct from the mouse *Apoe* gene, and the genes have species-specific promoters (Laws et al., 2003; Maloney et al., 2007). Second, mouse knock-in models have systemic alterations in ApoE production, while our manipulation was targeted to a specific brain region (the FC). Third, the relationship between *APOE* polymorphisms and transcription levels is unclear, making it unlikely that the direct manipulation of gene transcription in our study affects cellular function in the same way. Despite this, a foundational understanding of how *Apoe* functions in brain circuits that are implicated in cognitive processes will be critical for future research that targets this gene in disorders that feature attentional deficits, such as AD, schizophrenia, and attentional-deficit hyperactivity disorder (ADHD). The results of the current study highlight the need to study *Apoe,* attention, and anxiety in both biological sexes, and lay the groundwork for future studies on the mechanisms by which *Apoe* affects circuit function in the context of complex behavior.

## Materials and Methods

### Subjects

We used a cohort of 28 wild-type c57bl/6j mice (Jackson strain 000664) for all experiments. Of these mice, 14 were female (7 in the experimental Apoe group, 7 in the control EGFP group), and 14 were male (7 in the experimental Apoe group, 7 in the control EGFP group). Mice were group-housed (3-5 animals per cage) with *ad libitum* access to water. The colony room was temperature and humidity controlled on a 12 h light/dark cycle. All experiments were performed during the light cycle. At time of surgery, animals were roughly 90 to 120 days of age. All procedures were in accordance with the Institutional Animal Care and Use Committee of Lafayette College.

### Surgical Procedures

For all experiments, mice were anesthetized with isoflurane (1-2.5% oxygen) and then placed into a stereotaxic frame (Kopf Instruments, Tujunga, CA). An incision was made along the midline of the scalp, the skull was leveled, and bregma was identified. Holes were drilled in the skull above the FC (+1.7 mm AP from bregma, ±0.3 mm ML from the midline; Paxinos & Franklin, 2019), and an automated infusion pump (World Precision Instruments, Sarasota, FL) was used to inject the viruses at 4 nl/sec for a total volume of 600 nl/hemisphere (1.7 mm ventral to the surface of the brain). Experimental Apoe mice received injections of a virus coding for an Apoe expression cassette (AAV8-CMV-Apoe-EGFP; VectorBuilder custom catalog number AAV8-SP(VB900131-8416njv)-K0), and control EGFP mice received injections of a virus coding for EGFP alone (AAV8-CMV-EGFP; Addgene catalog # 105530). Following surgeries, mice were given 5 days to recover before handling.

### rCPT Training and Testing

Mice underwent five days of handling by the experimenter (10 minutes per day), and were introduced to the strawberry milkshake reward (Ensure Plus High Protein Nutrition Shake) in the home cage and placed on a food deprivation schedule (3 g of food per day) for the remainder of the experiment in order to maintain them at 90% of their *ad libitum* feeding weight. Following handling, mice were habituated to the testing chamber (Saksida-Bussey touchscreen chamber, catalog # 80614, Lafayette Instruments, Lafayette, IN) for two consecutive days (20 minutes per day) to ensure that they consumed the strawberry milkshake reward that was deposited in the reward trough at the rear of the chamber (Fig. 1a). Following habituation, mice were shaped to respond (physically touch the touchscreen at the front of the chamber) to a white square stimulus (stage 1; Fig. 1c), which was displayed for a 10 s period, or until the mouse touched the screen, at which point the stimulus disappeared. Successfully touching the screen when the white square was present (a “hit”) resulted in the presentation of a 1 s tone (3 kHz), reward trough illumination, and 20 μl of strawberry milkshake reward in the reward trough. The next trial was initiated when the mouse triggered the infra-red (IR) beam located in the reward trough. Stage 1 sessions were capped at 30 minutes. Once performance criteria were reached on stage 1 (>= 40 hits for two consecutive sessions), mice were moved to stage 2. During stage 2 sessions, the “target” image, or S+ (either vertical or horizontal bars, pseudo-randomized across mice so that an equal number of mice received both) was displayed on the touchscreen (2 s stimulus duration), and mice again were required to physically touch the screen when the target was displayed in order to get a reward. Performance criteria for stage 2 sessions (30 minutes in duration) were again >= 40 hits for two consecutive sessions.

Once mice reached performance criteria on stage 2, they were moved to stage 3 training. During stage 3 training, mice were shaped to respond (physically touch the screen) to the presentation of the S+, and withhold responding to another “non-target”, or S-, image (a snowflake). Stimulus duration was again 2 s. Mice were rewarded with strawberry milkshake for responding to the S+ (a “hit”), and were punished for responding to the S-(a “false alarm”) with a 20 second timeout period. Between stimulus presentations, mice waited for a variable intertrial interval (ITI) period of 2-3 s (randomly chosen). Mice could make four possible responses during stage 3 sessions: A hit (responding during an S+ presentation), a false alarm (responding during an S-presentation), a correct rejection (withholding responding during an S-presentation), and a miss (withholding responding during an S+ presentation). We used these response types to calculate a hit rate (defined as the number of hits divided by the total number of S+ presentations), a false alarm rate (defined as the number of false alarms divided by the total number of S-presentations), and a discrimination index, or d’ score (defined as the difference between the z-scored hit rate and false alarm rate) for each session. We also measured the hit latency (amount of time between stimulus presentation and response for hits), false alarm latency (amount of time between stimulus presentation and response for false alarms), and reward latency (amount of time between response and reward trough IR beam break for hits) for all sessions. Once mice reached a d’ score >= 0.6 for two consecutive sessions, they were given a final stage 3 probe session on the next day, which functioned similarly to other stage 3 training sessions with the exception of an altered probability of S+ presentation (20%, compared to 50% for other stage 3 training sessions). The probe session was designed to increase the attentional demand of the task (Methot and Huitema, 1998). Response types and latencies were collected with automated software (ABET II, Lafayette Instruments, Lafayette, IN). The chambers were cleaned thoroughly with 50% ethanol between sessions. Male and female mice were run in separate chambers throughout the duration of the experiment.

### Open Field Test

Following successful completion of the rCPT probe session, mice were placed into an open field apparatus (square enclosure measuring 60 cm x 60 cm, with 11 cm walls) In order to measure anxiety-like behavior and locomotion. The animals’ position within the apparatus was tracked at 30 fps, and the animals’ coordinates were recorded for each frame using ANY-maze tracking software (Stoelting, Wood Dale, IL). Each mouse was recorded in the open field for 10 minutes. ANY-maze software was also used to identify total time spent in the center of the apparatus, number of entries into the center, time spent freezing (lack of movement for >1 s), and total distance traveled throughout the 10 minute session. Open field testing was performed at least 1 day, and as long as 2 weeks, following rCPT completion.

### Single-Molecule Fluorescent In Situ Hybridization

In order to verify levels of *Apoe* transcription in FC tissue, we performed single-molecule *in situ* hybridization (RNAscope). Between 1 day and 1 week following open field testing, we killed mice via rapid cervical dislocation, extracted their brains, flash-froze them in 2-methylbutane, and stored them at −80 degrees C. We then took coronal sections of the FC (16 μm) on a cryostat (Leica, Nussloch, Germany), mounted them onto slides, and performed the RNAscope protocol using the fluorescent multiplex V2 kit from ACDBio (catalog # 323110). Specifically, tissue sections were briefly fixed with a 10% neutral buffered formalin solution at room temperature, and subjected to serious dehydration with ethanol. We then pretreated the sections with protease IV and hydrogen peroxide, and incubated the slides at 40 degrees C with a combination of three probes, which were as follows - channel 1: *Fos*, channel 2: *Gfp*, channel 3: *Apoe*. After incubation, we applied amplification buffers for each channel, and opal dyes (520 nm for channel 1; 570 nm for channel 2; 690 nm for channel 3; Akoya Biosciences, catalog #’s FP1487001KT, FP1488001KT, and FP1497001KT) in order to fluorescently label each transcript. Lastly, we stained the sections with DAPI to demarcate the nuclei of the cells. We then took z-stacked images of the FC (four sections per slide, one image per section, four images per mouse total) using a Zeiss LSM800 confocal microscope. For analysis, we used a MATLAB program to quantify transcript expression in each image (*dotdotdot*; Maynard et al., 2020). Specifically, we used the ‘CellSegm’ toolbox to perform nuclear segmentation in x, y, and z-dimensions to define regions of interest (ROIs) based on DAPI expression, and watershed analysis to identify distinct transcripts (individual dots) in each of the three microscope channels corresponding to an opal dye. We then co-localized each identified transcript with an identified nucleus (ROI). Transcripts that were not classified by the program as being co-localized with a nucleus were not used for analysis. Background noise, which could result from bleed-through from adjacent wavelengths in the gene channels, was eliminated using the ‘imhmin’ function. This function suppresses all of the minima in the grayscale image whose depth is less than the standard deviation of the image. We used a cutoff of 5 transcripts to categorize a cell as *Fos*-expressing (*Fos*+ cells), or *Gfp*-expressing (*Gfp*+).

### Statistical Analysis

For the number of sessions needed to reach criteria, open field results, and RNAscope summary statistics, we used two-way analyses of variance (ANOVAs) to compare each dependent variable between sex (male vs. female) and group (Apoe vs. EGFP). For rCPT performance metrics (latencies, hit rate, false alarm rate, d’, and c score), we used mixed-factorial ANOVAs (group x session) to compare dependent variables within sex. We used separate ANOVAs to compare these variables across the first five stage 3 training sessions (learning), and again between the final stage 3 session and the probe session (asymptotic performance). To analyze whether individual differences in *Apoe* expression were correlated with *Fos* expression and open field behavior, we performed linear regressions between these variables, and separated these correlations by sex, as our observed behavioral effects on the rCPT and open field tests differed by sex. Due to sample sizes that were too small to reliably observe possible interactions between these correlations, we used Cohen’s q and Fisher’s z-tests to calculate the difference in correlations between sexes (Cohen, 2016). All statistical tests were performed with the R programming language using RStudio (‘aov’ function for ANOVAs, ‘ggscatter’ function for linear regressions, and the ‘diffcor’ package for Cohen’s q and Fisher’s z-tests).

## Acknowledgements

The authors gratefully acknowledge Amy Badillo for animal care, and Jessie Greenlee for advice on statistical analysis. HLH also acknowledges support from a Brain and Behavior Research Foundation Young Investigator Award, and NIMH R21MH130066.

## Notes

### Competing Interest Statement

The authors have declared no competing interest.

